# Virulence studies of the human gut pathobiont *Bilophila wadsworthia* using *Galleria mellonella* as model host

**DOI:** 10.64898/2026.03.24.714029

**Authors:** Sara Matos, Beatriz Moniz, Dalila Mil-Homens, Inês A. C. Pereira, Andreia I. Pimenta

## Abstract

*Bilophila wadsworthia* is a gut pathobiont implicated in dysbiosis-driven inflammation, yet its pathogenic mechanisms remain poorly investigated. Here, we evaluated the suitability of *Galleria mellonella* larvae as an *in vivo* model to study *B. wadsworthia* infection. Two infection routes were compared: oral inoculation to mimic gastrointestinal colonization and hemolymph injection to model systemic infection. Oral challenge had minimal impact on larval health, whereas hemolymph injection caused marked morbidity, including reduced mobility, impaired cocoon formation, and progressive melanization, indicating that access to the circulatory system is required for overt disease. Infection required live bacteria, with *B. wadsworthia* capable of intracellular replication within hemocytes, leading to transient depletion of circulating immune cells followed by compensatory hemocyte proliferation. These findings reveal tight coupling between bacterial proliferation and host immune dynamics. Comparison with other sulfidogenic bacteria suggests that *Bilophila* pathogenicity is likely to involve host-specific interactions. Overall, our results establish *G. mellonella* as a practical and ethically favorable model to investigate *B. wadsworthia* virulence, host-pathogen interactions, and mechanisms relevant to gut-associated infection.

## 1. Introduction

The human gut microbiome is a complex and dynamic ecosystem, comprising trillions of microorganisms, including bacteria, viruses and fungi ^1^. This intricate community plays a crucial role in human health, influencing various physiological processes such as digestion, metabolism and immune function ^2,3^. The gut microbiome aids in breaking down dietary fibers, synthesizing essential vitamins, and protecting against pathogenic invaders ^4–6^. Dysbiosis of the human gut microbiota refers to an imbalance or maladaptation of the microbial communities within the gastrointestinal tract. This disruption can lead to a reduction in microbial diversity, an overgrowth of pathogenic species, and a decline in beneficial microbes, all of which can contribute to the development of various diseases, ranging from inflammatory gastrointestinal disorders to systemic conditions such as hypertension, obesity, diabetes, and even central nervous system-related disorders^7–13^. Factors contributing to dysbiosis include dietary changes, smoking, antibiotic usage, infections and chronic stress ^14–21^.

The Western diet, characterized by low-fiber, high-fat and high animal protein, significantly affects the stability of the gut microbial community ^22–24^. A high-fat diet is specifically associated with alterations in bile acid secretion and metabolism, with saturated fats promoting the taurine conjugation of hepatic bile acids ^25–27^. Elevated concentrations of such emulsifying conjugated bile acids increase intestinal taurine availability, promoting the expansion of taurine-metabolizing bacteria, sulfide-producing bacteria (sufidogenic bacteria, SB) such as *Bilophila wadsworthia* ^25,28–31^. *B. wadsworthia* is a Gram-negative, anaerobic bacterium that has attracted considerable interest in recent years owing to its association with various pathological conditions, particularly inflammatory disorders, colorectal cancer and metabolic diseases ^25,29,32,33^. This organism uses organic sulfonates, such as dietary or host-derived taurine and isethionate, as terminal electron acceptors, generating substantial amounts of hydrogen sulfide (H_2_S) through the dissimilatory sulfite reduction (Dsr) pathway^34–37^. Above physiological concentrations H_2_S is highly genotoxic, cytotoxic and interferes with the integrity of the intestinal epithelium and mucus barrier, inducing inflammation^38–40^. The association between *B. wadsworthia* and disease, particularly in the context of gut dysbiosis, has been largely attributed to the excessive production of H_2_S ^9,27,41–43^. Despite its low relative abundance in the healthy human microbiota, accounting for less than 0.01% of the normal flora, *B. wadsworthia* exhibits a high prevalence among humans ^44^. Additionally, it stands as one of the most frequently isolated microorganisms from anaerobic clinical infections, having been recovered from diverse extraintestinal sites, including the liver, bone, appendix, vagina, brain, and blood (bacteremia), suggesting a strong pathogenic potential^44–46^. However, the mechanisms underlying its pathogenicity and virulence traits remain largely unknown. Previous reports suggest the presence of hallmark virulence features in *B. wadsworthia* that are characteristic of Gram-negative pathogens. This bacterium exhibits the capacity to adhere to host cells^47^ and extracellular matrix proteins^48^, induce inflammation through the release of its lipopolysaccharide ^49^ and compromise the integrity of the colonic mucus layer ^40^.

Several studies using mice models have investigated the role of *B. wadsworthia* in disease^9,25,29,50^. Using immunocompromised mice, Devkota, et al. (2012) demonstrated that a high-fat diet promotes the expansion of *B. wadsworthia* and contributes to the development of inflammatory conditions ^25^. Additional studies have reported associations between intestinal colonization by *B. wadsworthia* and the development of metabolic dysfunction, including non-alcoholic fatty liver disease, obesity and chronic inflammatory diseases ^29,50^. More recently, Olson et al (2021) used various wild-type, germ-free and knock-out mouse models to establish a link between diet and hypoxia-related *B. wadsworthia* overgrowth, cognitive impairment and altered hippocampal physiology ^9^.

Given the capacity of *Bilophila* to trigger or enhance symptomatic diseases beyond the direct physiological effects of H_2_S, we sought to evaluate its virulence and pathogenic potential using a non-vertebrate infection system, *Galleria mellonella*. Commonly known as the greater wax moth, *G. mellonella* has emerged as a valuable and versatile model organism in microbiology, immunology, and toxicology research^51–56^. This insect model offers several advantages over traditional mammalian systems, including ease of handling, ethical feasibility, and a conserved innate immune response, making it an increasingly attractive platform for studying the virulence and host-pathogen interactions of diverse bacteria, fungal and yeast species ^52,54,56–61^. The use of *G. mellonella* circumvents many of the ethical concerns associated with vertebrate models. As invertebrates, these larvae are exempt from the ethical regulations governing mammalian research, allowing for greater flexibility in study design and enabling large scale, high-throughput analysis ^56,62,63^. *G. mellonella* can be maintained across a wide temperature range, including 37°C, the physiological human temperature ^64,65^, making it suitable to investigate pathogens under conditions that closely approximate those in the human host. *G. mellonella* is a cost-effective model, easy to maintain, and does not require specialized infrastructure, while its short life cycle and high reproductive rate facilitate rapid, reproducible experimentation. These features enable real-time monitoring of key parameters such as survival, immune response and microbial burden ^56,66^. Despite being an invertebrate, *G. mellonella* innate immune system shares significant functional and structural similarities with that of mammals ^53,66^. Its immune response comprises both cellular and humoral components, including phagocytosis and the production of antimicrobial peptides, making it a relevant model for studying infection dynamics and host-pathogen interactions^54,66–70^. Nonetheless, its inability to mount complex adaptive immune responses limits its application for investigating chronic or adaptive immunity-dependent diseases ^56,66,71^.

In this work, *G. mellonella* was used as experimental infection model to evaluate multiple biomarkers of *B. wadsworthia*-induced disease. Bacterial suspensions were used to establish both colonization and systemic infection, after which larval health status and survival were monitored. Viable bacterial load was quantified in the hemolymph, hemocytes were isolated and counted, and hemolymph melanization levels were determined. Overall, our data show that *G. mellonella* is a robust model for investigating *B. wadsworthia* pathogenicity and provide new insights into its underlying virulence mechanisms.

## 2. Methods

### 2.1 Bacterial strains

Four *B. wadsworthia* strains were used in this work: RZATAU strain (DSM 11045, DSMZ-German collection of microorganisms), ATCC 49260™ (American Type Culture Collection), and the human isolates QI0012 and QI0015 (kindly provided by Lizbeth Sayavedra and Arjan Narbad from the Quadram Institute Bioscience, UK) ^72^. All strains were grown and maintained in modified DSMZ 503 Anaerobic Freshwater (FWM) medium (MgCl_2_.6H_2_O (2 mM); NH_4_Cl (19 mM); CaCl_2_.2H_2_O (0.3 mM); KCl (7 mM); NaCl (17 mM); Na_2_SeO_3_ (11 µM); Na₂WO₄ (12 µM); trace element solution SL10 (1 mL/L); sodium resazurin (1.3 µM); 3-(N-morpholino)propanesulfonic acid - MOPS (50 mM); HCOONa (40 mM); Taurine 10 mM) and yeast extract (1 g/L)). The pH was corrected to 7.2, the solution dispensed into Hungate tubes and culture flasks and flushed with pressurized N_2_ before being sealed and sterilized by autoclaving. Before inoculation, the culture medium was supplemented with Thauer’s vitamin solution, Na_2_S (4 mM) and naftoquinone (200 µg/L). For CFU enumeration, *B. wadsworthia* was plated in 10 µL spots on BBE medium (Bacteroides Bile Esculin Agar Base, HiMedia Laboratories, #M805) and incubated anaerobically at 37°C until colonies could be counted.

*Nitratidesulfovibrio vulgaris* Hildenborough (DSM 644) was grown and maintained in DSMZ 63 *Desulfovibrio* Postgate medium at pH 7.0 and *N. vulgaris* Miyazaki F (DSM 19637) was grown and maintained in DSMZ 641 *Desulfovibrio* MV medium. All media were prepared following DSMZ instructions.

For infection assays, all bacterial strains were inoculated in fresh medium and incubated at 37°C without agitation. The optical density (OD) at 600 nm was monitored and bacterial cultures were grown until they reached the mid exponential phase (OD_600_ = 0.5-0.6).

### 2.2 Heat-Killed Bacterial Suspensions

For heat-killed (HK) inoculums, mid-exponential phase *B. wadsworthia* cultures were centrifuged at 4400 rpm for 15min. (Centrifuge 5702R, Eppendorf). In sterile and anaerobic conditions, the supernatant was discarded, and the bacterial pellet rinsed twice with PBS. The obtained pellet was resuspended with PBS to a final concentration of 2×10^8^ CFU/ml. The viable inoculum was incubated in a heated (80°C) bath for 1 h. As control, an aliquot of the same inoculum was maintained at room temperature. A 5 μL aliquot of each bacterial suspension (1×10^6^ CFU per larva) was injected into 10 larvae and, as control, a set of 10 larvae was injected with PBS. In all experiments, *inocula* were plated out for CFU count to validate the infectious dose and efficacy of the heat treatment.

### 2.3 Proteolytic Surface-Shaving

Mid-exponential phase bacterial cultures were collected as before. In sterile and anaerobic conditions, the supernatant was discarded, and the bacterial pellet rinsed twice with PBS. Final pellet was resuspended in PBS with 20% (w/v) of sucrose and the bacterial concentration was adjusted to obtain a final concentration of 2×10^8^ CFU/ml. Trypsin was added in a 2 µg per 1×10^7^ CFU proportion and the samples were then incubated at 37°C with mild agitation for 60 min. The untreated sample, with no trypsin, was also incubated at 37°C for 60 min. Following the incubation time, the bacterial suspensions were centrifuged for 2 min at 9000 rpm. Subsequently the pellet was suspended in the original volume of PBS to be used to inject *G. mellonella* larvae. In all experiments, *inocula* were plated to validate the infectious dose and bacterial viability after trypsin shaving.

### 2.4 B. wadsworthia lipopolysaccharide (LPS) extraction

*B. wadsworthia* strains and *N. vulgaris* Hildenborough (DSM 644) cells were grown and collected as previously described. LPS was extracted using LPS extraction kit (iNtRON) following the manufacturer instructions. The extracted LPS, resuspended in 10 mM Tris-HCl buffer (pH 8.0), was treated with 75 µg of protease K for 30 min at 50 °C, as suggested. The isolation and quality of the extracted LPS was assessed by a 16% Sodium dodecyl sulfate polyacrylamide gel electrophoresis (SDS-PAGE), followed by silver staining. To assess its virulent effect, 100 µg of LPS (in a 5 µL) was used to inject *G. mellonella* larvae.

### 2.5 G. mellonella growth and health monitorization

*Galleria mellonella* wax moth larvae were reared in our lab at 25 °C in the dark, from egg to last-instar larvae, on a natural diet (beeswax and pollen grains). Worms of the final-instar larval stage, weighing 250 ± 25 mg, were selected for the experiments.

For health monitorization, the incubated larvae were evaluated in periodic times of 24 h until 72 h, being given a score based on several health parameters such as movement, cocoon formation, melanization and survival (Table 1)^73^. The average punctuations for each parameter were then summed, giving a total score from 0 to 10 where 10 is the healthiest score and 0 the lowest. Infected larvae were considered dead when they failed to respond to physical stimulation. The results represent data from at least three independent experiments. For *G. mellonella* survival assay, 10 larvae were infected and larval survival was presented as a Kaplan-Meier survival curve, with data from at least three independent experiments.

**Table 1.** *G. mellonella* health index scoring system^73^.

| Category | Description | Score |
| --- | --- | --- |
| *Activity* | No movement | 0 |
|  | Minimal movement on stimulation | 1 |
|  | Move when stimulated | 2 |
|  | Move without stimulation | 3 |
| *Cocoon formation* | No cocoon | 0 |
|  | Partial cocoon | 0.5 |
|  | Full cocoon | 1 |
| *Melanization* | Black larvae | 0 |
|  | Black spots on brown larvae | 1 |
|  | ≥3 spots on beige larvae | 2 |
|  | <3 spots on beige larvae | 3 |
|  | No melanization | 4 |
| *Survival* | Dead | 0 |
|  | Alive | 2 |

### 2.6 G. mellonella intrahaemocoelic injection

Mid-exponential phase bacterial cultures were pelleted at room temperature. Cells were rinsed twice with PBS (4 mM KH_2_PO_4_; 16 mM Na_2_HPO_4_; 115 mM NaCl), pH 7.4, and then suspended in the same buffer. Bacterial suspensions were adjusted to the desired CFU inoculum by dilution. All bacterial manipulations were performed in anaerobic conditions.

Hemolymph injections were performed into the larvae last right proleg, previously sanitized with 70% (v/v) ethanol, using a micrometer-adapted microsyringe to control the volume of injection. Ten *G. mellonella* individuals were injected with 5 µL of different inoculum concentrations prepared in PBS and 10 exemplars with 5 µL of PBS as a mock control. For *B. wadsworthia* RZATAU strain the doses used were: 1×10^7^, 5×10^6^ and 1×10^6^ CFU per larva; for *Bilophila* ATCC, QI012 and QI015 strains 1×10^6^ CFU per larva was used, as well as for *N. vulgaris* strains. The larvae were then incubated at 37°C in absence of light for three days and the health status was monitored every 24 h. The bacterial load used for each assay was confirmed by plating the bacterial suspensions and CFU enumeration.

### 2.7 G. mellonella force-feeding infection

Force-feeding experiments involved groups of ten *G. mellonella* larvae, starved for 24 h prior to the test. The larvae were manually handled so that the needle fixed in the injector enters their mouth for force feeding of 5 µL of *B. wadsworthia* suspensions or PBS as mock control, using a micro syringe supported in a micrometer. Doses used were: 2×10^7^, 1×10^7^, 1×10^6^ and 1×10^5^ CFU per larva. The larvae were then incubated as described above.

### 2.8 Germ-free larvae

To assess the effect of the microbiota, germ-free *G. mellonella* were prepared. Briefly, larvae were treated by injection into the haemocoel, 24 h before the force-feeding assay, with two doses of 5 µL separated by 8h, with an antibiotic mix: Ampicillin (2 mg/mL), Kanamycin (2 mg/mL), Polymyxin B (2 mg/mL), Neomycin (2 mg/mL) and Vancomycin (1 mg/mL)^74^. Larvae were also treated in the same manner with PBS as control. Antibiotic and PBS treated larvae were then force-fed with a 1×10^7^ CFU inoculum per larvae and monitored as described before for four days (Figure 1).

**Figure 1.**
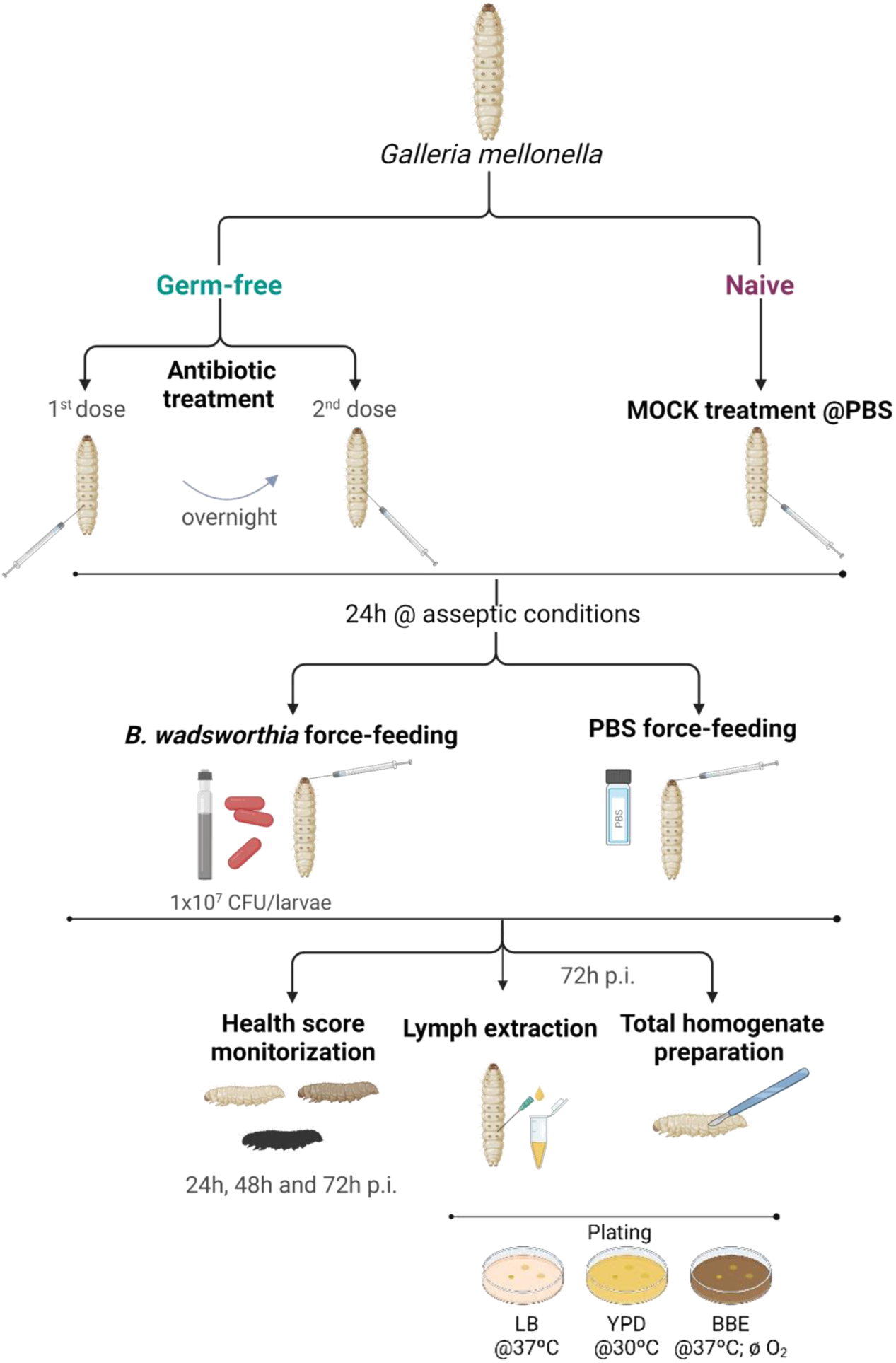
Schematic representation of the protocol for the germ-free larvae preparation.

To assess the bacterial colonization, in the end of the assay a pool of three larvae from each condition was macerated and a total homogenate was prepared. Larvae were collected, sanitized with 70% (v/v) ethanol and frozen at -20°C for 30 min. Them, using a homogenizer, the larvae were macerated together, and the resulting sample was diluted in Insect Physiological Saline solution (IPS) (150 mM NaCl; 5 mM KCl; 10 mM Tris pH 6.9; 10 mM EDTA; 30 mM sodium citrate) and plated in BBE plates under anaerobic conditions. To confirm the effectiveness of the antibiotic treatment, the obtained samples were also plated in Luria Broth (LB) and Yeast extract Peptone Dextrose (YPD) plates to determine bacterial and yeast growth, respectively. To assess the bacterial translocation from the larvae gastrointestinal tract to the hemocoel, hemolymph was extracted by piercing the area between the head and the thorax of three previously sanitized larvae, collected from each condition. The pooled sample was diluted in IPS buffer and plated in BBE, LB and YPD plates.

### 2.9 G. mellonella hemolymph collection

Groups of 20 larvae were inoculated with 1×10^6^ *B. wadsworthia* cells and the control larvae mock-infected with PBS. At 2, 6, 18, 24, 48 and 64 h after injection, three larvae were collected and sanitized with 70% (v/v) ethanol and hemolymph was extracted by piercing the area between the head and the thorax. Approximately, 50 µL of lymph were collected from each larva into a collection tube in sterilized conditions. The collected sample was used for assessment of *B. wadsworthia in vivo* proliferation, total hemocyte count, and melanization level.

### 2.10 Total hemocyte count, quantification of larvae melanization and measurement of bacterial proliferation

The collected hemolymph, from both *Bilophila*-infected and PBS mock-infected larvae, was diluted 1:10 in IPS. The total number of hemocytes was counted in a 10 µL sample using a cell counter (CellDrop™, DeNovix). To monitor melanization, absorbance was measured at 405 nm in a microplate reader (Lange DR 3900, Hach). IPS absorbance readings were used as a blank and the acquired values were subtracted from the sample readings. For measure of bacterial proliferation serial dilutions of lymph from both *Bilophila*-infected and PBS mock-infected larvae were performed in anaerobic conditions and plated in BBE medium. The plates were incubated at 37°C in sealed containers in anaerobic conditions until colonies were visible to count.

### 2.11 In vitro cultivation of hemocytes of G. mellonella

*G. mellonella* hemocytes were isolated as described previously^75^. Briefly, hemolymph was collected from larvae, and the outflowing hemolymph was immediately transferred into a sterile microtube containing IPS buffer in a 1:1 proportion. The hemolymph was centrifuged at 250xg for 10 min at 4°C to pellet hemocytes. The supernatant was discarded, and the pellet was washed twice with IPS and centrifuged at 250xg for 5 min at 4°C. The hemocyte pellet was then suspended gently in 1 ml of Grace insect medium (GIM) (Sigma) supplemented with 10% fetal bovine serum, 1% glutamine, and 1% antibiotic and antimycotic solution (10,000 U penicillin G, 10 mg streptomycin, 25 mg/liter amphotericin B). Suspended hemocytes were counted with a hemocytometer and incubated at 25°C in 24-well plates at a concentration of 2×10^5^ cells/well. Monolayers of primary *Galleria* hemocytes were used for experiments the next day.

### 2.12 Determination of bacterial load in hemocytes

*Galleria* hemocyte monolayer medium was replaced with GIM without antibiotics and oxygen, and then cells were infected with the bacterial suspensions with a MOI (multiplicity of infection) of 1:100. After 1 h of infection at 37°C under anaerobic conditions, the hemocytes were carefully washed twice with cell culture medium, followed by the addition of GIM containing 100 mg/L of gentamicin to kill the extracellular bacteria. After 1 h, supernatants were discarded, and the medium was replaced with GIM containing 25 mg/L of gentamicin. The quantification of viable intracellular bacteria was performed 2, 6, 8, 16 and 24 h after infection. Cell monolayers were lysed with 0.5% Triton X-100, and CFU were determined by plating dilutions of cell lysates on BBE-agar plates followed by incubation at 37°C under anaerobic conditions.

## 3 Results

### 3.1 *B. wadsworthia* causes symptomatic disease and eventually death in *G. mellonella*

To assess the consequences of *B. wadsworthia* systemic infection on the health and survival of *G. mellonella* larvae, *B. wadsworthia* cells were introduced directly into the lymphatic system via intrahaemocoelic injection. Alternatively, to evaluate the effects of gut colonization, which more accurately mimics the natural environment of *B. wadsworthia* in humans, the bacteria were introduced into the gastrointestinal system of larvae through force-feeding. In both scenarios, the health status of the larvae was monitored over three consecutive days and assessed using the standardized health index scoring system that considers movement, cocoon formation, melanization and survival ^73^ (Table 1, Figure 2A and 2B). Different bacterial loads were used for the two experiments: injection (1×10^6^, 5×10^6^ and 1×10^7^ CFU/larvae) and force-feeding (1×10^6^, 5×10^6^, 1×10^7^ and 2×10^7^ CFU/larvae). Health score assessments indicated that force-feeding did not result in a general decline in the larvae’s health status compared to the mock control, which was force-fed with PBS (Figure 2B). A significant decrease in health was only observed at the higher bacterial loads (1×10^7^ and 2×10^7^ CFU/larvae) with scores of 8.0 and 6.6 at 48 h and 8.8 and 7.9 at 72 h post-infection, respectively (P < 0.05 and < 0.01). In contrast, for systemic infection via injection all tested bacterial doses led to a significant reduction in health status (P < 0.0001) at all time points (Figure 2A). The data suggest that health status declines in a time-and dose-dependent manner, with higher bacterial load causing more severe health deterioration, and a progressive decline in health over time.

**Figure 2.**
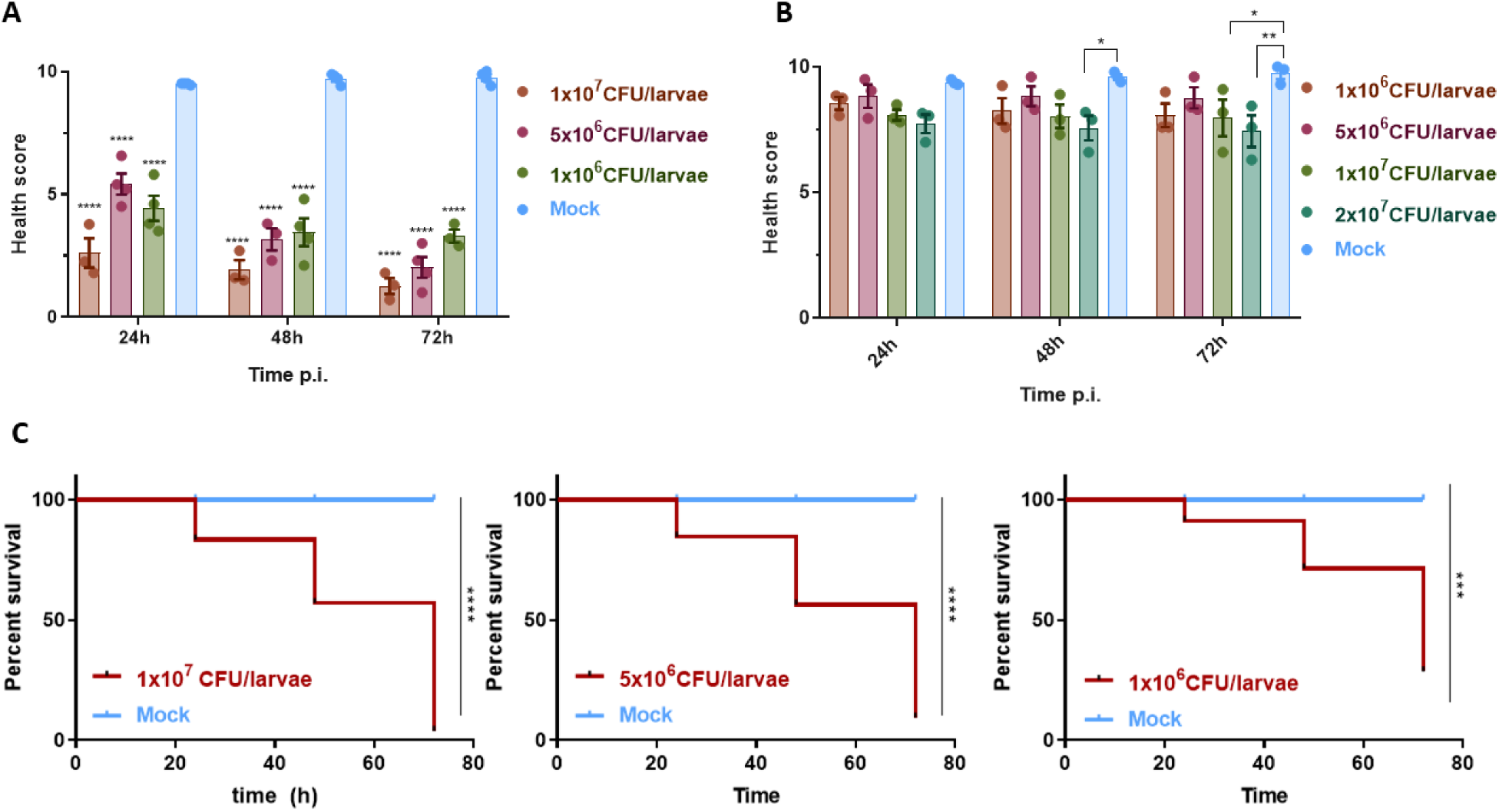
Health assessment and survival of *G. mellonella* wax worm larvae following *B. wadsworthia* challenge. (A) Post-infection health score of larvae injected with different loads of *B. wadsworthia*. (B) Post-infection health score of larvae force fed with different loads of *B. wadsworthia*. (C) Kaplan-Meier survival curves of infected wax worms infected. Mock – larvae injected only with PBS. (*P < 0.05; **P<0.01; P<0.0001)

To assess survival rates after infection, live and dead larvae were counted at each time point, and results represented using a Kaplan-Meier survival curve (Figure 2C). In the force-feeding experiment, the larvae survival rate remained at 100% across all time points (data not shown). In contrast, survival rates decreased in a dose-dependent manner for the injected larvae. At the highest dose (1×10^7^ CFU/larvae), the survival rate was 7%, while at the lowest dose (1×10^6^ CFU/larvae), the survival rate was 40% at 72 hours post-infection.

### 3.2 *B. wadsworthia* load is higher in the absence of *G. mellonella* native microbiota

To investigate the role of *G. mellonella* microbiota in regulating *B. wadsworthia* gut colonization, germ-free larvae were generated. These germ-free larvae were force-fed with 1×10^7^ CFU of *B. wadsworthia*, and their health scores compared to those of naive larvae infected with the same bacterial load (Figure 3). The health index score was assessed every 24 hours over four days. The results indicate that the presence or absence of the native microbiota does not affect the larvae’s response after *B. wadsworthia* inoculation, as in both cases there was no noticeable decline in health (Figure 3A), and no associated symptoms or mortality.

**Figure 3.**
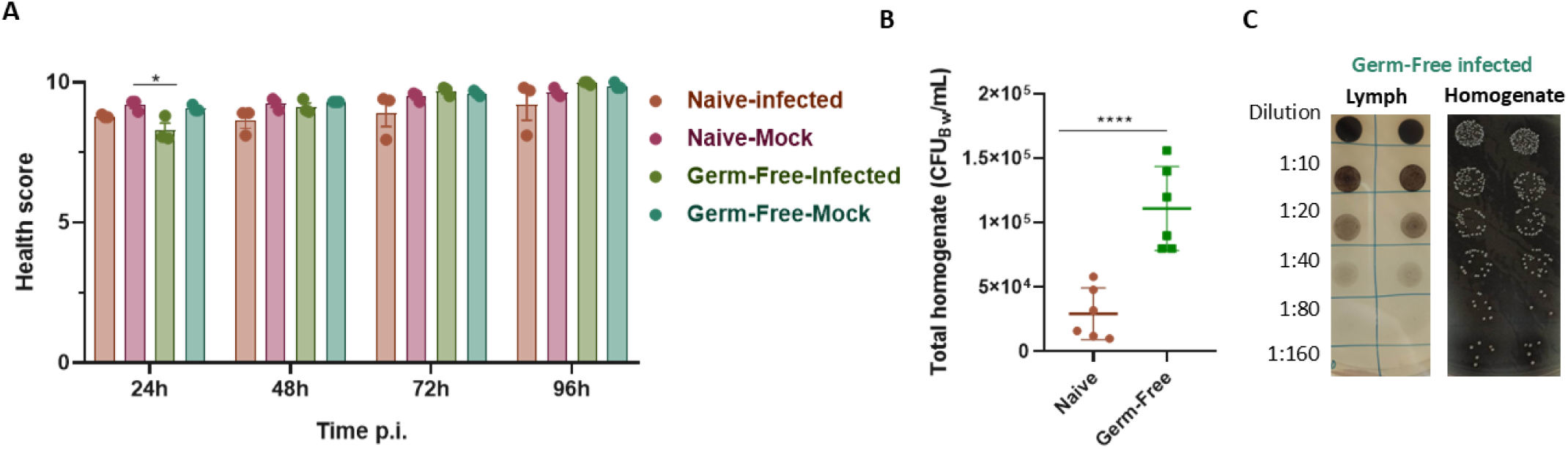
Health assessment and *B. wadsworthia* colonization of *G. mellonella* larvae. (A) Post-infection health score of the *G. mellonella* larvae force-fed with 1×10^7^ CFU of *B. wadsworthia*. (B) Load of *B. wadsworthia* (CFU/mL) collected form naïve- and germ-free-infected larvae. (C) BBE plates inoculated with haemolymph and total homogenate samples collected from germ free infected larvae. (*P < 0.05; **** P < 0.0001)

At the end of the four-day monitoring period, haemolymph was collected, and total homogenates were prepared from the larvae in each tested condition. No bacterial or yeast growth was detected in germ-free control larvae (force-fed with PBS), confirming that germ-free conditions were successfully maintained throughout the experiment. When comparing the *B. wadsworthia* load recovered from pooled homogenates of naïve-infected and germ-free infected larvae (Figure 3B), it was evident that the absence of the native microbiota led to significantly higher levels of *B. wadsworthia* in the gut of *G. mellonella* (P < 0.0001), confirming also the success of the force-feeding protocol. Additionally, no *B. wadsworthia* growth was detected in the haemolymph across all tested conditions (Figure 3C), in contrast to the homogenate samples where colonies are clearly seen and the agar plate turns black due to the production of sulfide. This indicates that there was no bacterial translocation from the gut to the circulatory system.

### 3.3 *B. wadsworthia* virulence is strain-specific and not general to all SB

To determine whether the effects of *B. wadsworthia* systemic infection are specific to this organism or are common to other sulfide-producing bacteria (sulfidogenic bacteria, SB), two *Nitratidesulfovibrio vulgaris* (previously *Desulfovibrio vulgaris*) strains were used to infect *G. mellonella* larvae (Figure 4A). *N. vulgaris* Hildenborough along with *N. vulgaris* Miyazaki F (NvM), were used to infect the wax worms with the same bacterial load as *B. wadsworthia* (1×10^6^ CFU/larvae). The larvae’s health was monitored every 24 hours for three consecutive days. The results, shown in Figure 4A, reveal that, unlike *B. wadsworthia* infection, larvae infected with *N. vulgaris* strains did not exhibit symptoms post-infection. The health scores of *Nitratiesulfovibrio*-infected larvae were nearly identical to those of the mock-infected larvae, and no mortality was observed. The same outcome was observed when higher bacterial loads (1×10^7^ and 2×10^7^ CFU/larvae) were tested, (data not shown). These findings suggest that the consequences of *G. mellonella* infection with *B. wadsworthia* are species specific.

**Figure 4.**
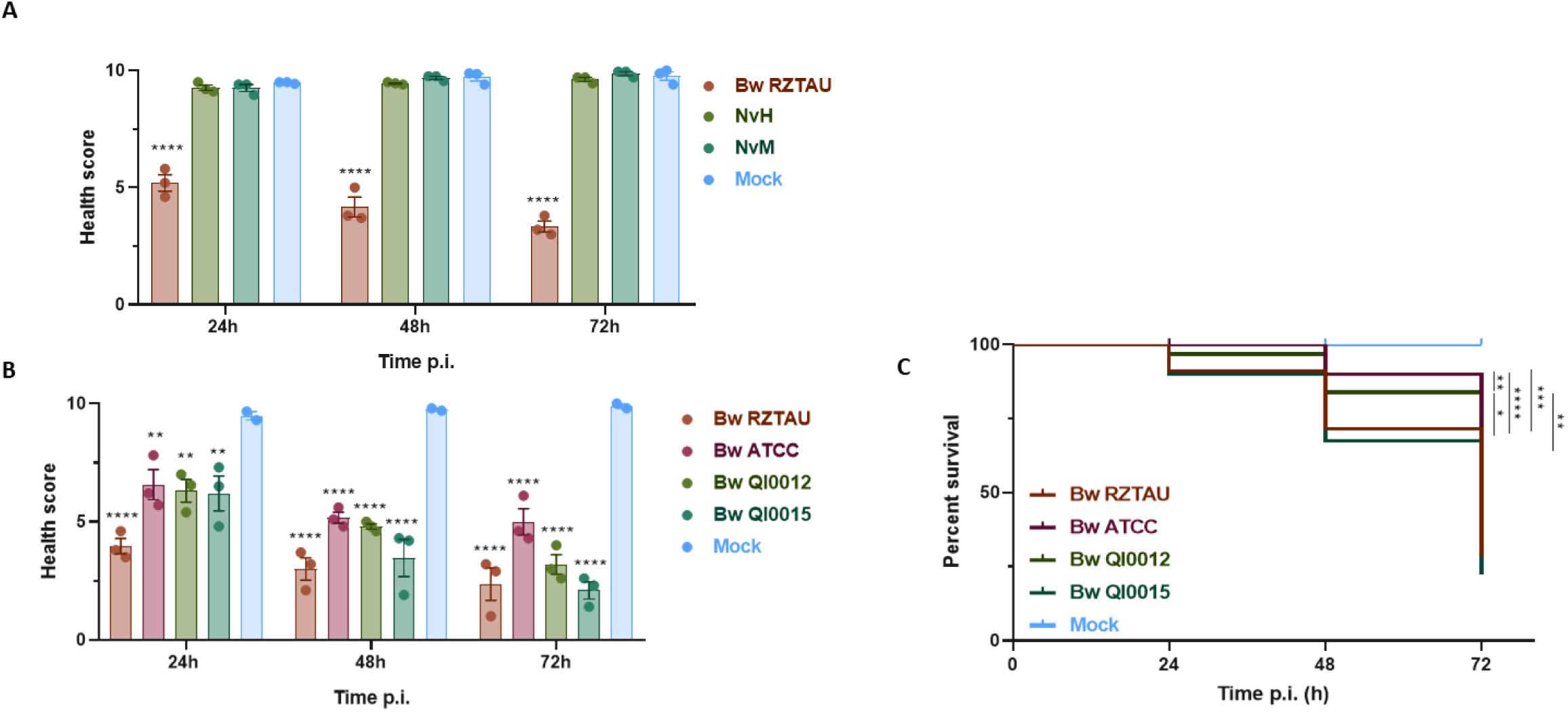
Specificity of *B. wadsworthia* virulence. (A) Post-infection health score of the *G. mellonella* larvae infected via injection with 1×10^6^ CFU/larvae of *B. wadsworthia* (Bw), *N. vulgaris* Hildenborough (NvH) and *N. vulgaris* Miyazaki F (NvM). (B) Post-infection health score of the *G. mellonella* larvae infected via injection with 1×10^6^ CFU/larvae of *B. wadsworthia* RZTAU (Bw RZTAU), *A*TCC 49260 (Bw ATCC), QI0012 (Bw QI0012) and QI0015 (Bw QI0015) strains. (C) Kaplan-Meier survival curves of infected larvae from B. Mock – larvae injected only with PBS. (* P<0.05; ** P<0.01; *** P<0.001; **** P<0.0001)

To evaluate the strain specificity of virulence, four different *B. wadsworthia* strains were tested: two laboratory strains—RZTAU (used in the previous assays) and ATCC 49260—and two strains isolated from stool samples of healthy human donors (QI0012 and QI0015)^72^. *G. mellonella* larvae were injected with equal bacterial loads (1×10^6^ CFU/larvae) of each strain, and the health index and survival rates were monitored (Figures 4B and 4C). All four strains caused a significant decline in larval health compared to the mock control (Figure 4B), but the degree and progression of disease varied. Based on both health scores and survival data, RZTAU and QI0015 emerged as the most virulent strains, inducing more severe health deterioration (Figure 4B) and lower survival rates at 72 h post-infection (29% and 23%, respectively; Figure 4C). In contrast to Bw RZTAU that causes symptomatic disease soon after 24 h post injection, Bw QI0015 needs more time to trigger an impairment on the larvae health. A similar behaviour occurs during infection with Bw QI0012, although the health score is a bit higher at all the time points. Regarding the percentage of survival, the results agree with the health scores, with Bw RZTAU and Bw QI0015 strains causing a more severe decline on *G. mellonella* survival rates at 72 h post infection - 29% and 23%, respectively. At this time, the percentage of survival for Bw ATCC and Bw QI0012 infection were 61% and 45%, indicating a lower virulence when compared to the two other strains.

### 3.4 *B. wadsworthia* infection is determined by bacterial viability

LPS, a major component of the outer membrane of Gram-negative bacteria, is a well-known immunostimulant and classical virulence factor. To assess its contribution to the strain specificity observed in *N. vulgaris* and *B. wadsworthia* infections, LPS was extracted from these organisms with a specific LPS extraction kit and evaluated for its ability to trigger an immune response and impair host health. LPS extracted from a normalized number of cells was purified and analyzed by 16% SDS-PAGE, showing minimal degradation (Figure S1). Low protein and nucleic acid contamination was also confirmed by spectrophotometric analysis. The extracted LPS was injected into the *G. mellonella* haemolymph, and larval health was monitored for three consecutive days (Figure 5A). No significant decrease (P < 0.05) in overall health was observed, regardless of LPS origin. Similarly, LPS injections into the haemolymph caused no mortality, with survival rates remaining 100% across all post-injection time points (data not shown). These findings indicate that LPS from both organisms does not trigger excessive activation of the *G. mellonella* innate immune system, as larval melanization remained unaffected throughout the assay. Together, these results suggest that *B. wadsworthia* employs virulent mechanisms other than LPS to impair host health and induce mortality during infection.

**Figure 5.**
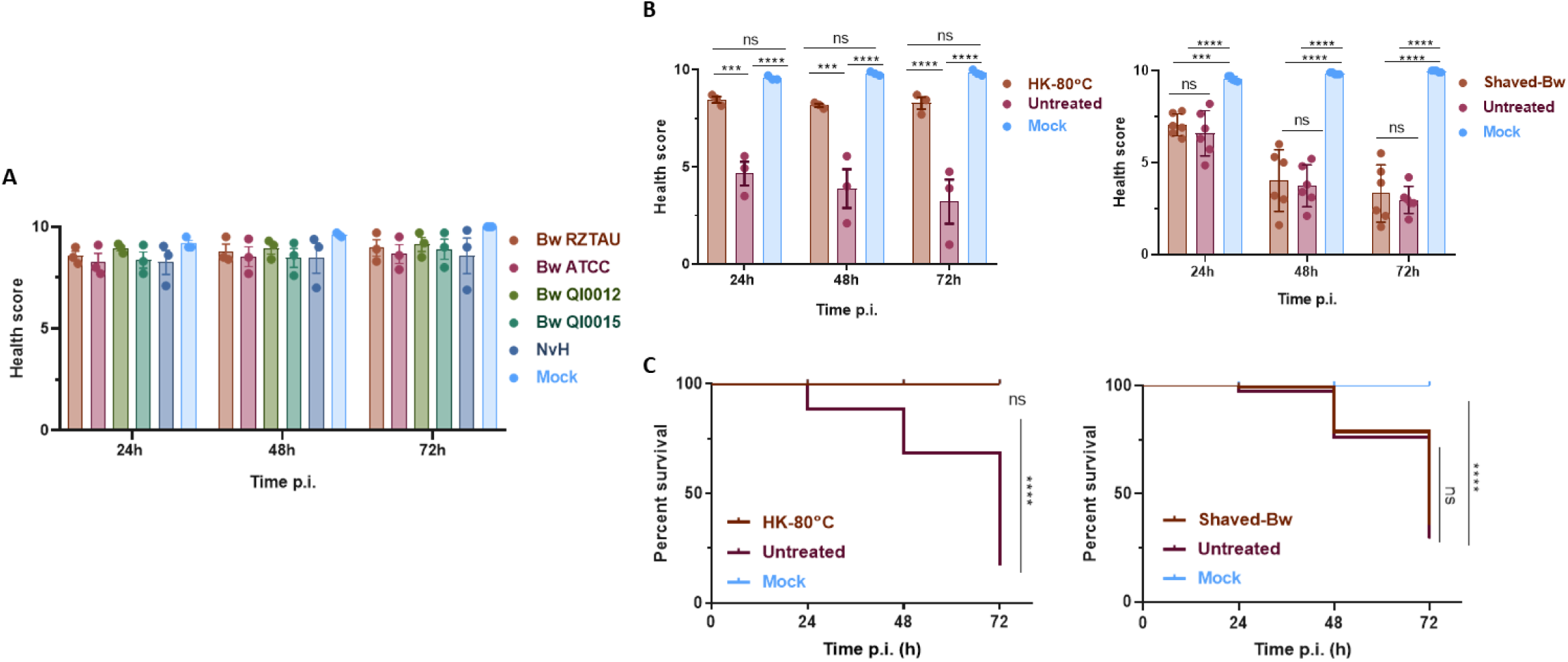
Study of *B. wadsworthia* virulence determinants. (A) Post-infection health score of *G. mellonella* larvae infected via injection with 100 µg of LPS extracted from *B. wadsworthia* RZTAU (Bw RZTAU), *A*TCC 49260 (Bw ATCC), QI0012 (Bw QI0012) and QI0015 (Bw QI0015) strains and *N. vulgaris* Hildenborough (NvH). (B) Health score of the *G. mellonella* larvae infected via injection with 1×10^6^ CFU/larvae of *B. wadsworthia* untreated, heat-killed at 80°C and after proteolytic shaving. (C) Kaplan-Meier survival curves of larvae infected via injection with 1×10^6^ CFU/larvae of *B. wadsworthia* untreated, heat-killed at 80°C and after proteolytic shaving. Mock – larvae injected only with PBS. (ns – nonsignificant (P > 0.05); *** P < 0.001; **** P < 0.0001)

To further assess the features of *B. wadsworthia* required to induce symptomatic disease, heat-killed bacteria and proteolytically shaved bacteria were used to infect *G. mellonella* larvae. Heat-killed bacteria were prepared by subjecting *B. wadsworthia* to a temperature of 80°C for 1 hour. The inactivation of *B. wadsworthia* cells was confirmed by the absence of growth on BBE plates inoculated with the heat-treated bacterial suspension. Proteolytic shaving of *B. wadsworthia* with trypsin for 1 h removed the bacterial surface protein appendages (confirmed by mass spectrometry) while maintaining cell viability (confirmed by plating and CFU counting). Both treated bacterial samples were injected into the *G. mellonella* haemolymph at the same bacterial load as the untreated bacteria, and the health score was assessed along the time (Figure 5B). The results indicate that active bacterial replication within the haemolymph is required for symptomatic disease, as heat-killed bacteria only caused a slight, non-significant (P > 0.05) decrease in the larvae’s health status, relative to the mock-infected controls, with 100% survival rate at all time points (Figure 5C). When the larvae were challenged with trypsin-shaved bacteria, their health condition remained very similar to that of larvae infected with untreated bacteria, with no significant differences (P > 0.05) (Figure 5B). The survival rates of larvae infected with shaved and unshaved *B. wadsworthia* were also very similar, falling below 50% at 72 h post-infection (Figure 5C). These results suggest that *B. wadsworthia* surface proteins are not required to induce *G. mellonella* death or health decline.

### 3.5 *B. wadsworthia* infection is characterized by bacterial proliferation and modulation of the larvae’s immune system

The impact of *B. wadsworthia* systemic infection on the physiology of *G. mellonella* larvae was assessed over the course of the infection. Haemolymph was collected from three larvae at each time point and pooled for analysis. This pooled sample was used to quantify melanization via absorbance (λ=405nm), count total haemocytes, and determine bacterial proliferation by counting *B. wadsworthia* CFUs (Figure 6). Melanization analysis revealed that melanin production was significantly higher at every time point compared to mock-infected larvae (P < 0.0001), and that it increased as the infection progressed, stabilizing after 48 h post-infection (Figure 6A). This increase in melanin production was visibly reflected in the progressive darkening of the larvae, as well as the colour of the collected haemolymph (Figure 6B).

**Figure 6.**
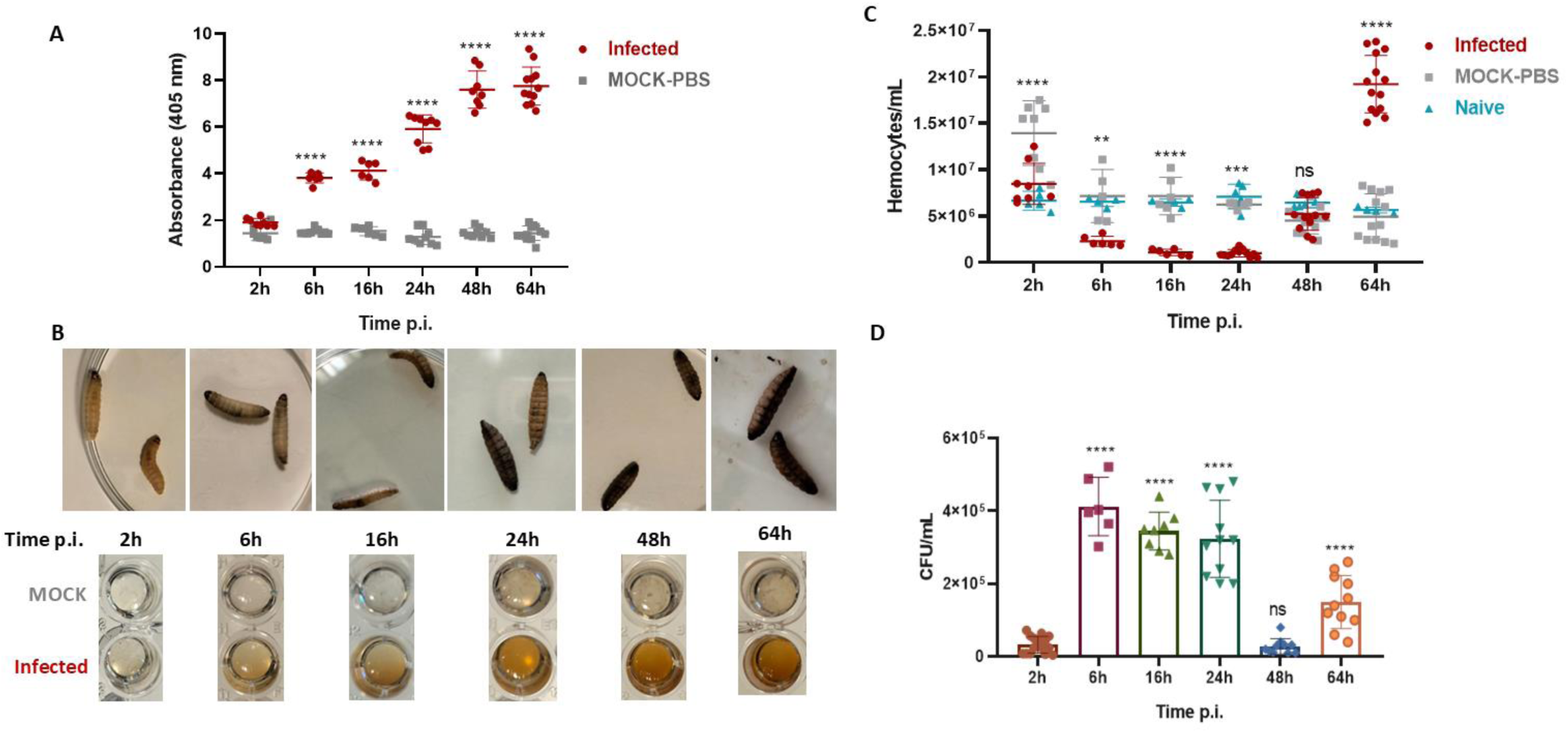
Key parameters of *B. wadsworthia* systemic infection in *G. mellonella*. (A) Post-infection haemolymph melanization from *B. wadsworthia*-infected (red) and mock- (grey) larvae. (B) Photographs of infected larvae and haemolymph collected over time of infection. (C) Total haemocyte count form naïve (blue), mock- (grey) and *B. wadsworthia* infected (red) larvae. (D) *B. wadsworthia* proliferation inside *G. mellonella* haemolymph during infection. Bacterial proliferation is represented as CFU/mL calculated from the growth observed after haemolymph plating. Statistical analyses were performed using the 2 h results as the reference for comparison. (ns – nonsignificant (P > 0.05); ** P < 0.01; *** P < 0.001; **** P < 0.0001)

Regarding total haemocyte counts, the results showed temporal variations (Figure 6C). At the initial time point (2 h), haemolymph from both *B. wadsworthia*-infected and mock-infected larvae exhibited a rise in haemocyte numbers compared to naïve larvae that had not undergone any puncture, suggesting it may be due to the physical damage caused by the injection. However, in *B. wadsworthia*-infected larvae, haemocyte numbers began to decline significantly by 6 h post-infection (P < 0.01), dropping approximately 8 times of the initial count by 24 h. At 48 h, the haemocyte count began to rise again, equalling the numbers of *B. wadsworthia*- and mock-infected larvae, and suggesting the onset of an immune response. By 64 h, the haemocyte count in *B. wadsworthia*-infected larvae was 2.4 times higher than at earliest time-point and 3.3 times higher than in mock-infected or naive larvae.

Bacterial proliferation within the haemolymph of infected larvae was determined by CFU counts (Figure 6D). The data demonstrate that *B. wadsworthia* cells are able to proliferate in the circulatory system of *G. mellonella* larvae. By 6 h post-infection, *B. wadsworthia* CFUs increased by 13-fold compared to the initial time point (2 h), a significant rise (P < 0.0001) that persisted until 24h post-infection. However, by 48 h, the bacterial load within the haemolymph of infected larvae had sharply decreased. Interestingly, with progression of infection, the bacterial load began to rise again, though at 64 h it did not reach the levels observed at 6-24 h post-infection. Haemolymph collected from non-infected larvae remained sterile, with no bacterial growth.

### 3.6 *B. wadsworthia* is able to invade and proliferate inside *G. mellonella* haemocytes

Given the observed variation in total haemocyte count and bacterial proliferation during infection, the ability of *B. wadsworthia* to invade, survive, and replicate within *G. mellonella* haemocytes was investigated. Haemolymph was collected from naïve larvae, and the haemocytes were isolated and cultured. These haemocytes were then challenged with *B. wadsworthia* at a MOI of 1:100, allowing the bacteria to enter the immune cells via phagocytosis. Following this, the culture supernatant, holding non-invading bacteria, was removed and replaced with fresh media supplemented with antibiotics for 1 h to ensure the complete elimination of extracellular bacteria (Figure 7). To assess the progression of infection, several time points were selected (2, 6, 8, 16, and 24 h), considering the natural viability of the haemocytes. At each time point, the haemocytes were lysed and the resulting suspension diluted and plated on BBE plates to quantify the bacterial load. The results showed a significant increase in the number of *B. wadsworthia* cells recovered from the cultured haemocytes over time (Figure 7). Intracellular replication was evident early in the infection, with bacterial concentrations increasing by 2.4 and 3.1 times at 6 and 8 h, respectively. A more pronounced and significant increase (11.4 times) was observed at 16 h post-infection, indicating robust bacterial replication within the haemocytes. This upward trend continued, with *B. wadsworthia* CFUs at 24 h post-infection being 14.1 times higher than at the initial time point (2 h), confirming the bacterium’s ability to effectively replicate inside *G. mellonella* haemocytes.

**Figure 7.**
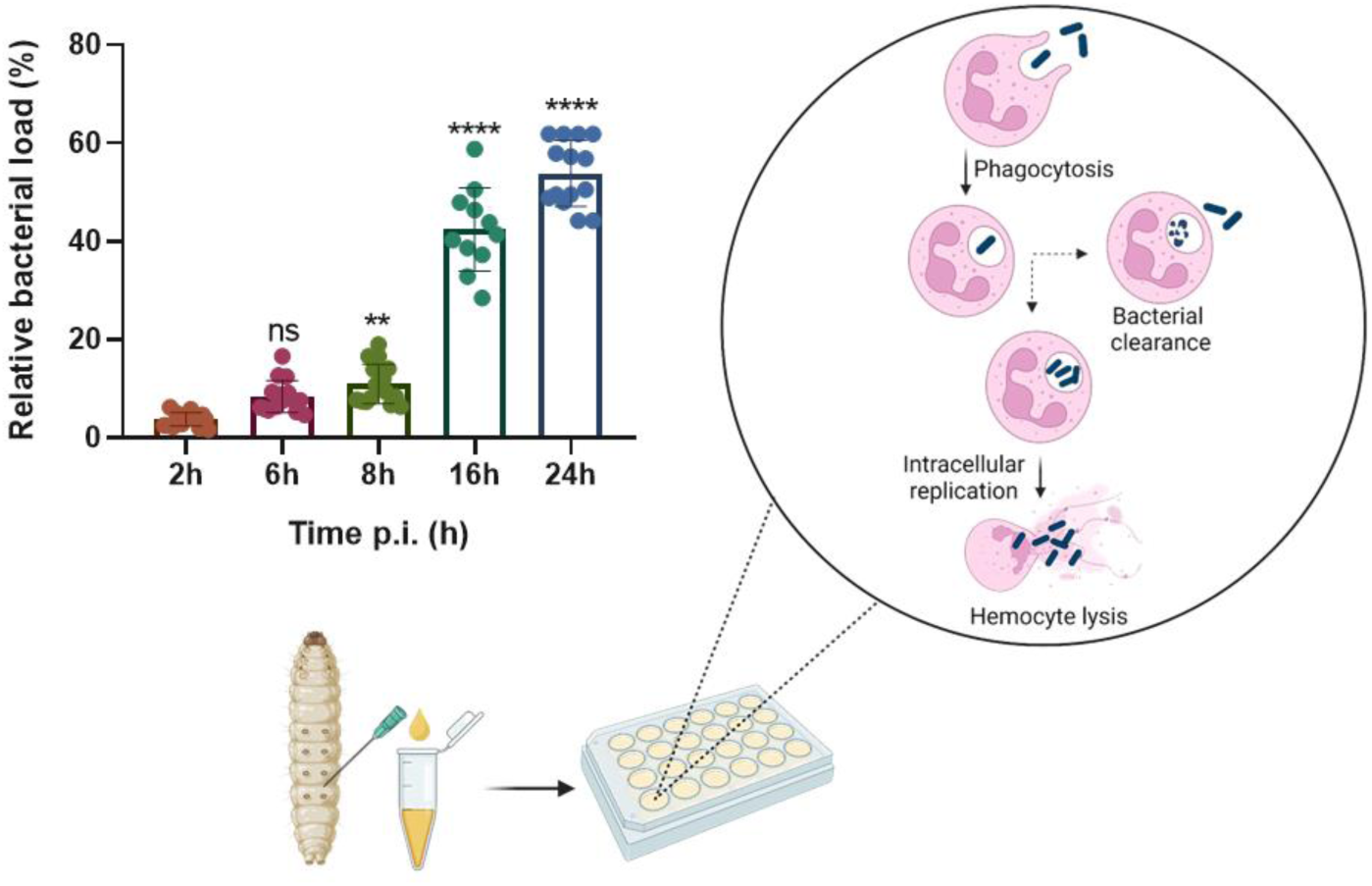
*B. wadsworthia* invasion and replication inside *G. mellonella* haemocytes. Schematic representation of the assay, focused on phagocytosis of *B. wadsworthia* by haemocytes and intracellular replication. Statistical analyses were performed using the 2 h results as the reference for comparison (ns – nonsignificant, **P<0.01, **** P<0.0001).

## 4 Discussion

The use of *G. mellonella* larvae as an *in vivo* infection model has become increasingly popular for studying a wide range of bacterial and fungal pathogens^54,55^. While this model has been extensively used to investigate various human pathogens, its application to the study of commensal or pathobiont members of the human gut microbiota remains limited. In our study, we provide evidence supporting the validity of this model for studying the infection and virulence mechanisms of the gut pathobiont *B. wadsworthia*.

We compared two routes of *B. wadsworthia* infection in *G. mellonella* mimicking gastrointestinal colonization (oral inoculation) and systemic infection (hemolymph injection). Injection caused severe pathology, whereas oral inoculation produced little to no effect on larval health. These findings suggest that *B. wadsworthia* requires translocation across the gastrointestinal epithelium to establish symptomatic infection. This is consistent with observations in humans, where *B. wadsworthia* is typically benign under normal colonization, but can promote epithelial damage and inflammation under dysbiotic conditions associated with overgrowth and elevated H_2_S production^27,41,42^, and also systemic dissemination leading to infection at several remote sites ^44–46^. In *G. mellonella*, however, *B. wadsworthia* was unable to cross the gut–hemocoel barrier. Even when colonization levels increased in the absence of the native microbiota, larval health remained unaffected, and no bacteria were recovered from the hemolymph. These results indicate that, in this model, direct entry into the circulatory system is required to elicit disease. In *Pseudomonas entomophila*, an insect gut pathogen, it has been shown that force-feeding infection requires a high bacterial load to kill larvae. Histological analyses revealed damage to the gut epithelium, including lysis and cellular degeneration, enabling dissemination into the hemocoel. This highlights the importance of pathogen–host specificity in triggering damage on the gut epithelium and breaching the gut barrier^76^. In contrast, *Mycobacterium abscessus*, *Staphylococcus aureus*, and *Pseudomonas aeruginosa* have been detected in gut tissues following hemocoel injection, likely due to passive dissemination from the surrounding fat body that encases the gut, rather than through an active invasion mechanism^77^.

Comparison of *B. wadsworthia* with other sulfidogenic bacteria, namely two *N. vulgaris* species, suggests that they may not share the same virulence mechanisms since these organisms had no adverse health effects. *B. wadsworthia* caused significantly more severe effects in the hemolymph infection model, suggesting that additional virulence traits contribute to disease in *G. mellonella*. These may include virulence factors that enable closer interaction with the host and that may be differentially expressed among strains, as suggested by the distinct pathogenic outcomes observed across *B. wadsworthia* isolates. These findings highlight the utility of the *G. mellonella* model in uncovering relevant virulence traits and support its application for robust species- and strain-level comparisons of pathogenicity.

Among the possible virulence factors identified for *B. wadsworthia*, LPS could be one of the most relevant, given its established role in the pathogenicity of many Gram-negative bacteria^49,78,79^. The virulent properties of *B. wadsworthia* LPS have been demonstrated *in vitro* using human mononuclear cells^49^, leading to abscess formation, although the effect is strongly influenced by the bacterial load and growth conditions^49^. In the *G. mellonella* model, however, injection with *B. wadsworthia* LPS did not cause significant alterations in larval health, in contrast to infection with whole bacteria. Given the well-known pro-inflammatory properties of Gram-negative LPS and its role in host immune modulation, an increase in melanization was expected for the LPS-infected larvae, since melanin production is a hallmark of the insect innate immune response^54^. While LPS from highly virulent *Pseudomonas aeruginosa* induced increased melanization, no effect was also observed with LPS isolated from several *Brucella* species, namely *B. abortus*, *B. melitensis*, and *B. microti*^80^. The ability of *Brucella* spp. to evade host immune recognition has been attributed to the low endotoxicity of their structurally modified LPS, which facilitates phagocytosis, resistance to intracellular killing, and replication before adaptive immunity is activated. A similar mechanism may underlie the weak immunostimulatory properties of *B. wadsworthia* LPS. Nonetheless, unlike *Brucella*^80^, injection of live *B. wadsworthia* into the *G. mellonella* hemolymph does lead to progressive melanization over the course of infection.

The ability of *B. wadsworthia* to adhere to host cells and extracellular matrix components has been previously reported^47,48^. This capacity is typically mediated by extracellular proteins, including adhesins and other surface-associated systems, which facilitate specific interactions between bacteria and host ligands^81,82^. To evaluate the contribution of such factors in our model, we employed an extracellular protein-shaving strategy to remove potential proteins involved in host interaction. Shaved and untreated bacteria were then injected into the hemolymph of *G. mellonella*, but no significant differences in host response were observed. These findings indicate that surface-associated proteins are not the key drivers of symptomatic infection in this invertebrate model. Nevertheless, extracellular proteins may still be important during the early stages of *B. wadsworthia*–host interactions in humans, where they may contribute to adhesion, invasion, toxin delivery, or immune modulation^81,83,84^. Thus, while dispensable for infection in *G. mellonella*, these proteins are likely to play a more prominent role within the natural niche of *B. wadsworthia*, the human gut.

Heat treatment, leading to impaired *B. wadsworthia* viability, resulted in a significant improvement in larval survival and health, compared with inoculation with untreated bacteria. These results indicate that *B. wadsworthia* viability and replicative capacity are crucial for establishing infection in *G. mellonella* following hemolymph inoculation. Loss of bacterial viability was associated with higher survival rates and improved health scores, suggesting that B*. wadsworthia* replication is required for the development of symptomatic disease in *G. mellonella*. This feature is not universal among Gram-negative pathogens. For example, in contrast to our findings, studies with *Escherichia coli* and *Klebsiella pneumoniae* have shown that heat-killed *inocula* injected into the hemocoel can still cause larval death, an effect attributed to the presence of dead bacterial cells rather than infection by live bacteria^85^. In the case of *B. wadsworthia*, bacterial viability may promote disease progression through the production of specific metabolites, in line with evidence that pathogens adapt their metabolism to exploit host-derived nutrients during infection^77^ and to modulate or evade immune stress responses^80^, or alter gene expression and protein production as part of adaptive responses to the host environment^86,87^.

To further investigate the role of bacterial viability in infection, we characterized *B. wadsworthia* proliferation over time and its role in modulating the host immune function. In response to a bacterial challenge, the innate immune system of *G. mellonella* engages both cellular and humoral components, in parallel to that of mammals. Hemocytes, the equivalent of mammalian white blood cells and central to the cellular immune response, comprise several subtypes with roles in phagocytosis, nodulation, and encapsulation^53,88^. Humoral immunity is triggered when pathogen-associated molecular patterns are recognized by host pattern recognition receptors, activating downstream signaling cascades. Among these, the prophenoloxidase (PPO) cascade drives melanization, leading to localized melanin deposition around invading microbes, a process functionally analogous to the mammalian complement cascade^89,90^. This melanin encapsulation is followed by hemocyte attachment and spreading, and ultimately by phagocytosis to clear the pathogen^89,90^. During infection, *B. wadsworthia* proliferated within the hemolymph with bacterial counts rising significantly up to 24 h post-infection. This is consistent with the requirement for viable bacteria to establish disease and reveals that the hemolymph environment supports bacterial replication. CFU counts dropped dramatically at 48 h before rising again at 64 h post-infection. This transient decline probably reflects the role of host immune defenses. To assess this, we monitored larval melanization and hemocyte counts across the same time points. Whereas melanization increased steadily throughout infection, hemocyte numbers did show a biphasic pattern, declining until 24 h and then recovering after 48 h. Comparison of bacterial proliferation and hemocyte dynamics suggests a close interplay between *B. wadsworthia* replication and temporary suppression or evasion of host immune responses, which fails to control the intense bacterial proliferation.

The correlation between reduced hemocyte numbers and rising bacterial loads has also been observed in other pathogens. *Listeria monocytogenes* infections in *G. mellonella* are marked by sustained bacterial proliferation from 4 to 24 h, accompanied by steadily increasing melanization and a 90% drop in viable hemocytes compared to PBS-injected controls at 24 h post-infection^91^. The authors suggest that the insect death could result from the significant reduction in hemocyte numbers caused directly by *L. monocytogenes*^91^. Similarly, *K. pneumoniae* infection leads to larval death through robust bacterial replication, heightened melanization (via PO activity), and hemocyte depletion^87^. Nevertheless, this pattern contrasts with *G. mellonella* infections by *Brucella* spp., where reduced immune recognition leads to stable CFU levels across 72 h, with the exception of *B. microti*, which displays an early decline in bacterial counts at 6 h post-infection, likely reflecting more efficient immune clearance^80^.

Given their phagocytic ability, isolated *Galleria* hemocytes were challenged with *B. wadsworthia ex vivo*. The results revealed that *Bilophila* is not only internalized by host cells, but it also replicates intracellularly, as the number of recovered CFUs increased more than 14-fold within 24 h post-infection. This strong intracellular proliferation likely leads to cell lysis, explaining the marked reduction in hemocyte counts observed *in vivo* up to 24 h post-infection. The subsequent increase in hemocyte numbers at 48 h likely represents a stress-driven host compensatory response, leading to the production of new immune cells to counteract the increased bacterial burden. This response also likely accounts for the transient drop in *B. wadsworthia* CFUs. However, despite the strong hemocyte increase until 64 h, this immune response was not sufficient to control a second bacterial expansion, suggesting that at that point the health damage was beyond recovery and bacteria overwhelmed the host, ultimately leading to larval death.

In similar studies with other pathogens, Torres et al. demonstrated that inhibiting phagocytosis or hemocyte increase enhances *G. mellonella* susceptibility to *E. coli* and *K. pneumoniae*, underscoring the importance of viable circulating hemocytes in controlling bacterial proliferation^85^. This suggests that there is an effective link between the number of viable circulating haemocytes and the capacity of bacterial pathogens to proliferate inside the larvae. For *Brucella*, Elizalde-Bielsa et al. showed that the number of infected hemocytes and intracellular bacteria slightly decreased 48 h post-infection, consistent with the low endotoxicity of *Brucella* LPS and the corresponding delayed mortality and limited melanization response^80^. In contrast, *K. pneumoniae* was not detected within hemocytes at 5, 12, or 24 h post-infection, indicating that *Galleria* hemocytes does not engulf this pathogen^87^. Instead, hemocyte numbers declined steadily up to 24 h, suggesting direct pathogen-induced immune cell destruction^87^. Taken together, these observations support the conclusion that larval mortality and melanization are directly linked to the degree of pathogen recognition and engagement by the host innate immune system.

In conclusion, our study establishes *G. mellonella* as a robust and informative *in vivo* model for investigating *B. wadsworthia* pathogenicity. We show that symptomatic infection requires bacterial access to the hemolymph, where live bacteria that replicate both extracellularly and within hemocytes trigger dynamic host immune responses, including melanization and changes in hemocyte population. The severity of infection, coupled with the requirement for bacterial viability and the inability of host defenses to fully control bacterial proliferation, highlights the multifactorial nature of *B. wadsworthia* virulence. Furthermore, comparison with other pathogens highlights the role of pathogen-specific interactions with the host innate immune system in shaping disease outcome. Overall, our findings validate *G. mellonella* as a practical and ethically favorable platform for dissecting gut pathobiont virulence mechanisms, enabling species- and strain-level studies of gut-associated bacterial infections.

## Acknowledgments

The authors would like to acknowledge Lizbeth Sayavedra and Arjan Narbad from the Quadram Institute Bioscience, Norwich, UK, for providing us with the QI0012 and QI0015 *Bilophila wadsworthia* strains.

## Authors’ contributions

AIP, DMH and IACP conceived and designed the project. AIP and DMH designed the experiments. SM, BM and AIP performed experiments. SM and BM contributed equally to the manuscript. AIP, SM, BM and IACP analyzed the data. AIP wrote the manuscript with contributions from all coauthors.

## References

1. Deschasaux M, Bouter KE, Prodan A, et al. Depicting the composition of gut microbiota in a population with varied ethnic origins but shared geography. Nat Med. 2018;24(10):1526–1531. doi:10.1038/s41591-018-0160-1

2. Guarner F, Malagelada J R. Gut flora in health and disease. Lancet. 2003;361(9356):512–519. doi:10.1016/S0140-6736(03)13438-1

3. Thursby E, Juge N. Introduction to the human gut microbiota. Biochem J. 2017;474(11):1823–1836. doi:10.1042/BCJ20160510

4. Zhang Y, Chen R, Zhang DD, Qi S, Liu Y. Metabolite interactions between host and microbiota during health and disease: Which feeds the other? Biomed Pharmacother. 2023;160(December 2022):114295. doi:10.1016/j.biopha.2023.114295

5. Postler TS, Ghosh S. Understanding the Holobiont: How Microbial Metabolites Affect Human Health and Shape the Immune System. Cell Metab. 2017;26(1):110–130. doi:10.1016/j.cmet.2017.05.008

6. Kamada N, Chen GY, Inohara N, Núñez G. Control of pathogens and pathobionts by the gut microbiota. Nat Immunol. 2013;14(7):685–690. doi:10.1038/ni.2608

7. Li J, Zhao F, Wang Y, et al. Gut microbiota dysbiosis contributes to the development of hypertension. Microbiome. 2017;5(1):1–19. doi:10.1186/s40168-016-0222-x

8. Sun MF, Shen YQ. Dysbiosis of gut microbiota and microbial metabolites in Parkinson’s Disease. Ageing Res Rev. 2018;45:53–61. doi:10.1016/j.arr.2018.04.004

9. Olson CA, Iñiguez AJ, Yang GE, et al. Alterations in the gut microbiota contribute to cognitive impairment induced by the ketogenic diet and hypoxia. Cell Host Microbe. 2021;29(9):1378–1392.e6. doi:10.1016/j.chom.2021.07.004

10. Rajca S, Grondin V, Louis E, et al. Alterations in the intestinal microbiome (Dysbiosis) as a predictor of relapse after infliximab withdrawal in Crohn’s disease. Inflamm Bowel Dis. 2014;20(6):978–986. doi:10.1097/MIB.0000000000000036

11. Duboc H, Rajca S, Rainteau D, et al. Connecting dysbiosis, bile-acid dysmetabolism and Gut inflammation in inflammatory bowel diseases. Gut. 2013;62(4):531–539. doi:10.1136/gutjnl-2012-302578

12. Halfvarson J, Brislawn CJ, Lamendella R, et al. Dynamics of the human gut microbiome in inflammatory bowel disease. Nat Microbiol. 2017;2(February):1–7. doi:10.1038/nmicrobiol.2017.4

13. Sommer F, Bäckhed F. The gut microbiota-masters of host development and physiology. Nat Rev Microbiol. 2013;11(4):227–238. doi:10.1038/nrmicro2974

14. Arumugam M, Raes J, Pelletier E, et al. Enterotypes of the human gut microbiome. Nature. 2011;473(7346):174–180. doi:10.1038/nature09944

15. Bai X, Wei H, Liu W, et al. Cigarette smoke promotes colorectal cancer through modulation of gut microbiota and related metabolites. Gut. 2022;71(12):2439–2450. doi:10.1136/gutjnl-2021-325021

16. Bernard-Raichon L, Venzon M, Klein J, et al. Gut microbiome dysbiosis in antibiotic-treated COVID-19 patients is associated with microbial translocation and bacteremia. Nat Commun. 2022;13(1):1–13. doi:10.1038/s41467-022-33395-6

17. Ding S, Chi MM, Scull BP, et al. High-fat diet: Bacteria interactions promote intestinal inflammation which precedes and correlates with obesity and insulin resistance in mouse. PLoS One. 2010;5(8). doi:10.1371/journal.pone.0012191

18. Fishbein SRS, Mahmud B, Dantas G. Antibiotic perturbations to the gut microbiome. Nat Rev Microbiol. 2023;21(12):772–788. doi:10.1038/s41579-023-00933-y

19. Lee K, Raguideau S, Sirén K, et al. Population-level impacts of antibiotic usage on the human gut microbiome. Nat Commun. 2023;14(1):1191. doi:10.1038/s41467-023-36633-7

20. Rodriguez J, Hiel S, Neyrinck AM, et al. Discovery of the gut microbial signature driving the efficacy of prebiotic intervention in obese patients. Gut. 2020;69(11):1975–1987. doi:10.1136/gutjnl-2019-319726

21. Zhang X, Coker OO, Chu ESH, et al. Dietary cholesterol drives fatty liver-associated liver cancer by modulating gut microbiota and metabolites. Gut. 2021;70(4):761–774. doi:10.1136/gutjnl-2019-319664

22. Li Z tao, Wang J wei, Hu X hai, et al. The effects of high-fat foods on gut microbiota and small molecule intestinal gases: release kinetics and distribution in vitro colon model. Heliyon. 2022;8(10):e10911. doi:10.1016/j.heliyon.2022.e10911

23. Rinninella E, Cintoni M, Raoul P, et al. Food components and dietary habits: Keys for a healthy gut microbiota composition. Nutrients. 2019;11(10):1–23. doi:10.3390/nu11102393

24. Ruengsomwong S, La-ongkham O, Jiang J, Wannissorn B, Nakayama J, Nitisinprasert S. Microbial Community of Healthy Thai Vegetarians and Non-Vegetarians, Their Core Gut Microbiota, and Pathogen Risk. J Microbiol Biotechnol. 2016;26(10):1723–1735. doi:10.4014/jmb.1603.03057

25. Devkota S, Wang Y, Musch MW, et al. Dietary-fat-induced taurocholic acid promotes pathobiont expansion and colitis in Il10-/- mice. Nature. 2012;487(7405):104–108. doi:10.1038/nature11225

26. Ou J, Carbonero F, Zoetendal EG, et al. Diet, microbiota, and microbial metabolites in colon cancer risk in rural Africans and African Americans. Am J Clin Nutr. 2013;98(1):111–120. doi:10.3945/ajcn.112.056689

27. Ridlon JM, Wolf PG, Gaskins HR. Taurocholic acid metabolism by gut microbes and colon cancer. Gut Microbes. 2016;7(3):201–215. doi:10.1080/19490976.2016.1150414

28. Devkota S, Chang EB. Interactions between diet, bile acid metabolism, gut microbiota, and inflammatory bowel diseases. Dig Dis. 2015;33(3):351–356. doi:10.1159/000371687

29. Natividad JM, Lamas B, Pham HP, et al. Bilophila wadsworthia aggravates high fat diet induced metabolic dysfunctions in mice. Nat Commun. 2018;9(1):2802. doi:10.1038/s41467-018-05249-7

30. Larabi AB, Masson HLP, Bäumler AJ. Bile acids as modulators of gut microbiota composition and function. Gut Microbes. 2023;15(1):1–27. doi:10.1080/19490976.2023.2172671

31. Alrehaili BD, Lee M, Takahashi S, et al. Bile acid conjugation deficiency causes hypercholanemia, hyperphagia, islet dysfunction, and gut dysbiosis in mice. Hepatol Commun. 2022;6(10):2765–2780. doi:10.1002/hep4.2041

32. Baron EJ, Summanen P, Downes J, Roberts MC, Wexler H, Finegold SM. Bilophila wadsworthia, gen. nov. and sp. nov., a Unique Gram-negative Anaerobic Rod Recovered from Appendicitis Specimens and Human Faeces. Microbiology. 1989;135(12):3405–3411. doi:10.1099/00221287-135-12-3405

33. Nguyen LH, Ma W, Wang DD, et al. Association Between Sulfur-Metabolizing Bacterial Communities in Stool and Risk of Distal Colorectal Cancer in Men. Gastroenterology. 2020;158(5):1313–1325. doi:10.1053/j.gastro.2019.12.029

34. Peck SC, Denger K, Burrichter A, Irwin SM, Balskus EP, Schleheck D. A glycyl radical enzyme enables hydrogen sulfide production by the human intestinal bacterium Bilophila wadsworthia. Proc Natl Acad Sci U S A. 2019;116(8):3171–3176. doi:10.1073/pnas.1815661116

35. Laue H, Friedrich M, Ruff J, Cook AM. Dissimilatory sulfite reductase (Desulfoviridin) of the taurine-degrading, non-sulfate-reducing bacterium Bilophila wadsworthia RZATAU contains a fused DsrB-DsrD subunit. J Bacteriol. 2001;183(5):1727–1733. doi:10.1128/JB.183.5.1727-1733.2001

36. Laue H, Denger K, Cook AM. Taurine reduction in anaerobic respiration of Bilophila wadsworthia RZATAU. Appl Environ Microbiol. 1997;63(5):2016–2021. doi:10.1128/aem.63.5.2016-2021.1997

37. da Silva SM, Venceslau SS, Fernandes CLV, Valente FMA, Pereira IAC. Hydrogen as an energy source for the human pathogen Bilophila wadsworthia. *Antonie van Leeuwenhoek*, Int J Gen Mol Microbiol. 2008;93(4):381–390. doi:10.1007/s10482-007-9215-x

38. Dordević D, Jančíková S, Vítězová M, Kushkevych I. Hydrogen sulfide toxicity in the gut environment: Meta-analysis of sulfate-reducing and lactic acid bacteria in inflammatory processes. J Adv Res. 2021;17(27):55–69. doi:10.1016/j.jare.2020.03.003

39. Attene-Ramos MS, Nava GM, Muellner MG, Wagner ED, Plewa MJ, Gaskins HR. DNA damage and toxicogenomic analyses of hydrogen sulfide in human intestinal epithelial FHs 74 Int cells. Environ Mol Mutagen. 2010;51(4):304–314. doi:10.1002/em.20546

40. Ijssennagger N, van der Meer R, van Mil SWC. Sulfide as a Mucus Barrier-Breaker in Inflammatory Bowel Disease? Trends Mol Med. 2016;22(3):190–199. doi:10.1016/j.molmed.2016.01.002

41. Pimenta AI, Bernardino RM, Pereira IAC. Role of sulfidogenic members of the gut microbiota in human disease. Adv Microb Physiol. 2024;85:145–200.

42. Carbonero F, Benefiel AC, Alizadeh-Ghamsari AH, Gaskins HR. Microbial pathways in colonic sulfur metabolism and links with health and disease. Front Physiol. 2012;3 NOV(November):1-11. doi:10.3389/fphys.2012.00448

43. Murros KE. Hydrogen Sulfide Produced by Gut Bacteria May Induce Parkinson’s Disease. Cells. 2022;11(6). doi:10.3390/cells11060978

44. Baron EJ, Curren M, Henderson G, et al. Bilophila wadsworthia isolates from clinical specimens. J Clin Microbiol. 1992;30(7):1882–1884. doi:10.1128/jcm.30.7.1882-1884.1992

45. Baron EJ. Bilophila wadsworthia: a Unique Gram-negative Anaerobic Rod. Anaerobe. 1997;3(2-3):83–86. doi:10.1006/anae.1997.0075

46. Nava GM, Carbonero F, Croix JA, Greenberg E, Gaskins HR. Abundance and diversity of mucosa-associated hydrogenotrophic microbes in the healthy human colon. ISME J. 2012;6(1):57–70. doi:10.1038/ismej.2011.90

47. Gerardo SH, Garcia MM, Wexler HM, Finegold SM, Gerardo SH. Adherence of Bilophila wadsworthia to cultured human embryonic intestinal cells. Anaerobe. 1998;4(1):19–27. doi:10.1006/anae.1997.0134

48. Schumacher UK. Adherence of Bilophila wadsworthia to Laminin and Fibronectin. Clin Infect Dis. 1997;25(s2):S180–S180. doi:10.1086/516180

49. Mosca A, D’Alagni M, del Prete R, et al. Preliminary evidence of endotoxic activity of Bilophila wadsworthia. Anaerobe. 1995;1(1):21–24. doi:10.1016/S1075-9964(95)80379-3

50. Henao-Mejia J, Elinav E, Jin C, et al. Inflammasome-mediated dysbiosis regulates progression of NAFLD and obesity. Nature. 2012;482(7384):179–185. doi:10.1038/nature10809

51. Moya-Andérico L, Vukomanovic M, Cendra M del M, et al. Utility of Galleria mellonella larvae for evaluating nanoparticle toxicology. Chemosphere. 2021;266. doi:10.1016/j.chemosphere.2020.129235

52. Curtis A, Binder U, Kavanagh K. Galleria mellonella Larvae as a Model for Investigating Fungal—Host Interactions. Front Fungal Biol. 2022;3(April):1–12. doi:10.3389/ffunb.2022.893494

53. Gallorini M, Marinacci B, Pellegrini B, et al. Immunophenotyping of hemocytes from infected Galleria mellonella larvae as an innovative tool for immune profiling, infection studies and drug screening. Sci Rep. 2024;14(1):759. doi:10.1038/s41598-024-51316-z

54. Smith DFQ, Dragotakes Q, Kulkarni M, Hardwick JM, Casadevall A. Galleria mellonella immune melanization is fungicidal during infection. Commun Biol. 2022;5(1):1–13. doi:10.1038/s42003-022-04340-6

55. Lange A, Schäfer A, Bender A, et al. Galleria mellonella: A novel invertebrate model to distinguish intestinal symbionts from pathobionts. Front Immunol. 2018;9(SEP):1–12. doi:10.3389/fimmu.2018.02114

56. Dinh H, Semenec L, Kumar SS, Short FL, Cain AK. Microbiology’s next top model: Galleria in the molecular age. Pathog Dis. 2021;79(2):ftab006. doi:10.1093/femspd/ftab006

57. Barnoy S, Gancz H, Zhu Y, Honnold CL, Zurawski D V., Venkatesan MM. The Galleria mellonella larvae as an in vivo model for evaluation of Shigella virulence. Gut Microbes. 2017;8(4):335–350. doi:10.1080/19490976.2017.1293225

58. Sänger PA, Wagner S, Liebler-Tenorio E, Fuchs TM. Dissecting the invasion of Galleria mellonella by Yersinia enterocolitica reveals metabolic adaptations and a role of a phage lysis cassette in insect killing. PLoS Pathog. 2022;18(11):1–26. doi:10.1371/journal.ppat.1010991

59. Mil-Homens D, Martins M, Barbosa J, et al. Carbapenem-resistant klebsiella pneumoniae clinical isolates: In vivo virulence assessment in galleria mellonella and potential therapeutics by polycationic oligoethyleneimine. Antibiotics. 2021;10(1):1–14. doi:10.3390/antibiotics10010056

60. Pais P, Vagueiro S, Mil-homens D, et al. A new regulator in the crossroads of oxidative stress resistance and virulence in Candida glabrata : The transcription factor CgTog1. Virulence. 2020;11(1):1522–1538. doi:10.1080/21505594.2020.1839231

61. Romera D, Aguilera-Correa JJ, Garciá-Coca M, et al. The Galleria mellonella infection model as a system to investigate the virulence of Candida auris strains. Pathog Dis. 2020;78(9):1–7. doi:10.1093/femspd/ftaa067

62. Trevijano-Contador N, Zaragoza O. Expanding the use of alternative models to investigate novel aspects of immunity to microbial pathogens. Virulence. 2014;5(4):454–456. doi:10.4161/viru.28775

63. Jander G, Rahme LG, Ausubel FM. Positive Correlation between Virulence of Pseudomonas aeruginosa Mutants in Mice and Insects. J Bacteriol. 2000;182(13):3843–3845. doi:10.1128/JB.182.13.3843-3845.2000

64. Réjasse A, Gilois N, Barbosa I, et al. Temperature-dependent production of various PlcR-controlled virulence factors in Bacillus weihenstephanensis strain KBAB4. Appl Environ Microbiol. 2012;78(8):2553–2561. doi:10.1128/AEM.07446-11

65. Mil-homens D, Barahona S, Moreira RN, et al. Stress Response Protein BolA Influences Fitness and Promotes Salmonella enterica Serovar Typhimurium Virulence. Appl Environ Microbiol. 2018;84(8):e02850–17.

66. Kavanagh K, Reeves EP. Exploiting the potential of insects for in vivo pathogenicity testing of microbial pathogens. FEMS Microbiol Rev. 2004;28(1):101–112. doi:10.1016/j.femsre.2003.09.002

67. Bergin D, Reeves EP, Renwick J, Wientjes FB, Kavanagh K. Superoxide production in Galleria mellonella hemocytes: Identification of proteins homologous to the NADPH oxidase complex of human neutrophils. Infect Immun. 2005;73(7):4161–4170. doi:10.1128/IAI.73.7.4161-4170.2005

68. Strand MR. Insect Hemocytes and Their Role in Immunity. Insect Immunol. 2008;32:25–47. doi:10.1016/B978-012373976-6.50004-5

69. Brown SE, Howard A, Kasprzak AB, Gordon KH, East PD. A peptidomics study reveals the impressive antimicrobial peptide arsenal of the wax moth Galleria mellonella. Insect Biochem Mol Biol. 2009;39(11):792–800. doi:10.1016/j.ibmb.2009.09.004

70. Cerenius L, Söderhäll K. The prophenoloxidase-activating system in invertebrates. Immunol Rev. 2004;198(1):116–126. doi:10.1111/j.0105-2896.2004.00116.x

71. Asai M, Li Y, Newton SM, Robertson BD, Langford PR. Galleria mellonella-intracellular bacteria pathogen infection models: The ins and outs. FEMS Microbiol Rev. 2023;47(2):1–32. doi:10.1093/femsre/fuad011

72. Davies J, Mayer MJ, Juge N, Narbad A, Sayavedra L. Bacteroides thetaiotaomicron enhances H2S production in Bilophila wadsworthia. Gut Microbes. 2024;16(1). doi:10.1080/19490976.2024.2431644

73. Loh JMS, Adenwalla N, Wiles S, Proft T. Galleria mellonella larvae as an infection model for group A streptococcus. Virulence. 2013;4(5):419–428. doi:10.4161/viru.24930

74. Kong HG, Kim HH, Chung J, et al. The Galleria mellonella Hologenome Supports Microbiota-Independent Metabolism of Long-Chain Hydrocarbon Beeswax. Cell Rep. 2019;26(9):2451–2464.e5. doi:10.1016/j.celrep.2019.02.018

75. Admella J, Torrents E. A Straightforward Method for the Isolation and Cultivation of Galleria mellonella Hemocytes. Int J Mol Sci. 2022;23(21):13483. doi:10.3390/ijms232113483

76. Kordaczuk J, Sułek M, Mak P, Śmiałek-Bartyzel J, Hułas-Stasiak M, Wojda I. Defence response of Galleria mellonella larvae to oral and intrahemocelic infection with Pseudomonas entomophila. Dev Comp Immunol. 2023;147(February 2023):104749. doi:10.1016/j.dci.2023.104749

77. Admella J, Torrents E. Investigating bacterial infections in Galleria mellonella larvae: Insights into pathogen dissemination and behavior. J Invertebr Pathol. 2023;200(May):107975. doi:10.1016/j.jip.2023.107975

78. Mohr AE, Crawford M, Jasbi P, Fessler S, Sweazea KL. Lipopolysaccharide and the gut microbiota: considering structural variation. FEBS Lett. 2022;596(7):849–875. doi:10.1002/1873-3468.14328

79. Freudenberg MA, Merlin T, Gumenscheimer M, Kalis C, Landmann R, Galanos C. Role of lipopolysaccharide susceptibility in the innate immune response to Salmonella typhimurium infection: LPS, a primary target for recognition of Gram-negative bacteria. Microbes Infect. 2001;3(14-15):1213–1222. doi:10.1016/S1286-4579(01)01481-2

80. Elizalde-Bielsa A, Aragón-Aranda B, Loperena-Barber M, et al. Development and evaluation of the Galleria mellonella (greater wax moth) infection model to study Brucella host-pathogen interaction. Microb Pathog. 2023;174(October 2022):2–9. doi:10.1016/j.micpath.2022.105930

81. Pimenta AI, Mil-homens D, Kilcoyne M, et al. Burkholderia cenocepacia BCAM2418-induced antibody inhibits bacterial adhesion, confers protection to infection and enables identification of host glycans as adhesin targets. Cell Microbiol. 2021;(e13340). doi:10.1111/cmi.13340

82. Pimenta AI, Mil-Homens D, Fialho AM. Burkholderia cenocepacia – host cell contact controls the transcription activity of the trimeric autotransporter adhesin. Microbiologyopen. 2020;9(e998). doi:10.1002/mbo3.998

83. Stones DH, Krachler AM. Against the tide: The role of bacterial Adhesion in host colonization. Biochem Soc Trans. 2016;44(6):1571–1580. doi:10.1042/BST20160186

84. Pizarro-Cerdá J, Cossart P. Bacterial adhesion and entry into host cells. Cell. 2006;124(4):715–727. doi:10.1016/j.cell.2006.02.012

85. Torres MP, Entwistle F, Coote PJ. Effective immunosuppression with dexamethasone phosphate in the Galleria mellonella larva infection model resulting in enhanced virulence of Escherichia coli and Klebsiella pneumoniae. Med Microbiol Immunol. 2016;205(4):333–343. doi:10.1007/s00430-016-0450-5

86. Bruchmann S, Feltwell T, Parkhill J, Short FL. Identifying virulence determinants of multidrug-resistant Klebsiella pneumoniae in Galleria mellonella. Pathog Dis. 2021;79(3):1–15. doi:10.1093/femspd/ftab009

87. Insua JL, Llobet E, Moranta D, et al. Modeling Klebsiella pneumoniae Pathogenesis by Infection of the Wax Moth Galleria mellonella. Bliska JB, ed. Infect Immun. 2013;81(10):3552–3565. doi:10.1128/IAI.00391-13

88. Browne N, Heelan M, Kavanagh K. An analysis of the structural and functional similarities of insect hemocytes and mammalian phagocytes. Virulence. 2013;4(7):597–603. doi:10.4161/viru.25906

89. Nakhleh J, El Moussawi L, Osta MA. The Melanization Response in Insect Immunity. In: Advances in Insect Physiology. Vol 52. 1st ed. Elsevier Ltd.; 2017:83–109. doi:10.1016/bs.aiip.2016.11.002

90. González-Santoyo I, Córdoba-Aguilar A. Phenoloxidase: A key component of the insect immune system. Entomol Exp Appl. 2012;142(1):1–16. doi:10.1111/j.1570-7458.2011.01187.x

91. Joyce SA, Gahan CGM. Molecular pathogenesis of Listeria monocytogenes in the alternative model host Galleria mellonella. Microbiology. 2010;156(11):3456–3468. doi:10.1099/mic.0.040782-0

