## Supplementary material for "Virulence studies of the human gut pathobiont *Bilophila wadsworthia* using *Galleria mellonella* as model host": Figure S1

**Figure S1-** Sodium dodecyl sulfate polyacrylamide gel electrophoresis (SDS-PAGE) of

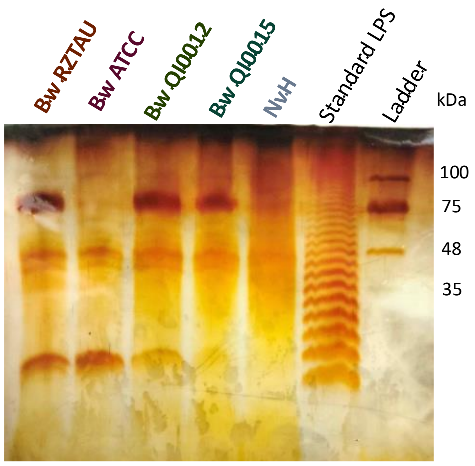


isolated LPS from *B. wadsworthia* RZTAU, ATCC, QI0012, QI0015 and *N. vulgaris* Hildenborough wt strains. The isolation and quality of the extracted LPS was assessed by a 16% SDS-PAGE followed by silver staining. Marker - NZYColour Protein Marker II. Standard LPS -Lipopolysaccharides from *Salmonella enterica* serotype typhimurium (Sigma-Aldrich).
